# Development of Anti-Inflammatory Extracellular Vesicles by Surface Expression of Syndecan-4

**DOI:** 10.1101/2025.07.05.663259

**Authors:** Lijuan Yu, Markus Bergqvist, Kyong-Su Park, Cecilia Lässer, Jan Lötvall

## Abstract

The biological functions of extracellular vesicles (EVs) depend on their cellular source. Further, different subpopulations of EVs from the same cells carry different cargo, but differences in their biological functions are less understood. We here identify a very small EV subpopulation released by HEK293F cells (miniEVs). These EVs, in contrast to the larger EVs, were found to have anti-inflammatory properties. Quantitative proteomics identified a potential anti-inflammatory molecule, Syndecan-4 (SDC4), on the surface of the miniEVs, but not larger EVs. We engineered HEK293F cells to overexpress SDC4, which results in the molecule being highly expressed in all EV subpopulations. Expression of SDC4, a proteoglycan, also increased the presence of heparan sulfate on the EV surface. Furthermore, these EVs were found to have potent anti-inflammatory effects *in vitro*, which heparinase treatment could slightly reduce. Furthermore, the SDC4 EVs showed anti-inflammatory effects *in vivo* in a model of peritonitis. We conclude that HEK293F EVs can be engineered to become anti-inflammatory, and that SDC4-expressing HEK293F-EV potentially could become an anti-inflammatory therapeutic.

## Introduction

Inflammation is a fundamental defence mechanism that protects the body from harmful stimuli [1]. However, excessive inflammation, triggered by infection or severe trauma, can be harmful [2]. Glucocorticoids have been the cornerstone of anti-inflammatory therapy for over 50 years, and while effective, their use is associated with significant side effects, such as osteoporosis [3, 4]. In recent decades, monoclonal antibodies targeting specific inflammatory molecules, such as IL-4, IL-5, IL-6, IL-13, and TNF-α, have emerged as treatments for specific diseases [5]. However, these therapies lack the broad anti-inflammatory effects of glucocorticoids, and are weak against innate immune responses mediated by broad Toll-like receptor (TLR) activation, such as those seen in cytokine storms or sepsis [6]. Consequently, there remains a critical need for novel therapies broadly targeting inflammation, particularly in conditions where current treatments fall short [7].

Cell-based therapies, particularly those involving mesenchymal stem cells (MSCs), have shown promise in treating degenerative and inflammatory diseases [8–10]. Notably, the FDA recently approved an MSC-based therapy for steroid-refractory acute graft-versus-host disease (GvHD) in children, underscoring the potential of such approaches [11]. However, the use of live cell therapies poses challenges related to manufacturing, storage, and distribution [12, 13]. Importantly, research has revealed that the therapeutic effects of MSCs are partly mediated by their secretome, particularly extracellular vesicles (EVs) [14]. EVs are lipid-bilayer enclosed nanoparticles released by many different cell types. They carry a multitude of membrane lipids and membrane proteins, as well as cargo RNA and intravesicular proteins. When derived from MSC, EVs have demonstrated significant anti-inflammatory potential, offering a promising cell-free alternative to traditional cell therapies [15]. Two double-blind, placebo-controlled studies have demonstrated that MSC-EVs accelerated recovery and showed trends toward improved survival in severe inflammatory diseases [16, 17]. A third study found that topical application of MSC-EVs enhanced skin healing after laser therapy compared to placebo [18]. Nonetheless, challenges such as low EVs production yield, limited MSCs passages, and rapid cellular senescence pose significant obstacles to the development of MSC-EV-based anti-inflammatory therapies.

An alternative to developing MSC-EVs for the purpose of creating anti-inflammatory therapeutics is to engineer HEK293F cells to produce EVs with anti-inflammatory properties. For example, engineering the anti-inflammatory molecule CD24 into HEK293F EVs has been performed, and these EVs have been clinically tested in cytokine storms associated with COVID-19 [19]. In brief, inhalation of CD-24 EVs resulted in rapid resolution of pulmonary inflammation, and patients could be released from hospital care earlier than without that treatment.

We hypothesized that different subpopulations of EVs released by cells may convey different biological functions. To test this, we isolated different subpopulations of EVs from HEK293F cells, which is “Large EVs” and “Small EVs” isolated by differential ultracentrifugation methods [20], and explored whether the remaining secretome contains even smaller EV components. We explored whether any unique features in the EVs were associated with the expression of different proteins, which was explored using quantitative proteomics. Candidate anti-inflammatory molecules were overexpressed in EVs by genetic engineering of HEK293F cells, after which their functions were explored *in vitro* and *in vivo*.

## Results

### Isolation and characterization of three subpopulations of HEK293F EVs

We first isolated large and small EVs (hereafter referred to as L-EVs and S-EVs) from HEK293F cells, using differential ultracentrifugation at 16,500 × g (L-EVs) and 118,000 × g (S-EVs) [21]. We then added an additional step of 118,000 × g for 16h to also isolate particles with smaller sizes that had not been pelleted in the two previous steps (hereafter referred to as miniEVs). All EV pellets were further purified on iodixanol density cushions (Extended data Figure 1a). Transmission electron microscopy (TEM) showed that all three EV subtypes have round morphology (Figure 1a and Extended Figure 1b). Manual measurements of single-EV diameters using TEM revealed that the three EV subtypes have different average sizes (Figure 1b). Next, the purity of the three subpopulations before and after the iodixanol purification process was calculated as particle-to-protein ratios (Figure 1c-e). Notably, HEK293F S-EVs and miniEVs exhibited improved purity after iodixanol density gradient centrifugation [21].

**Figure 1.**
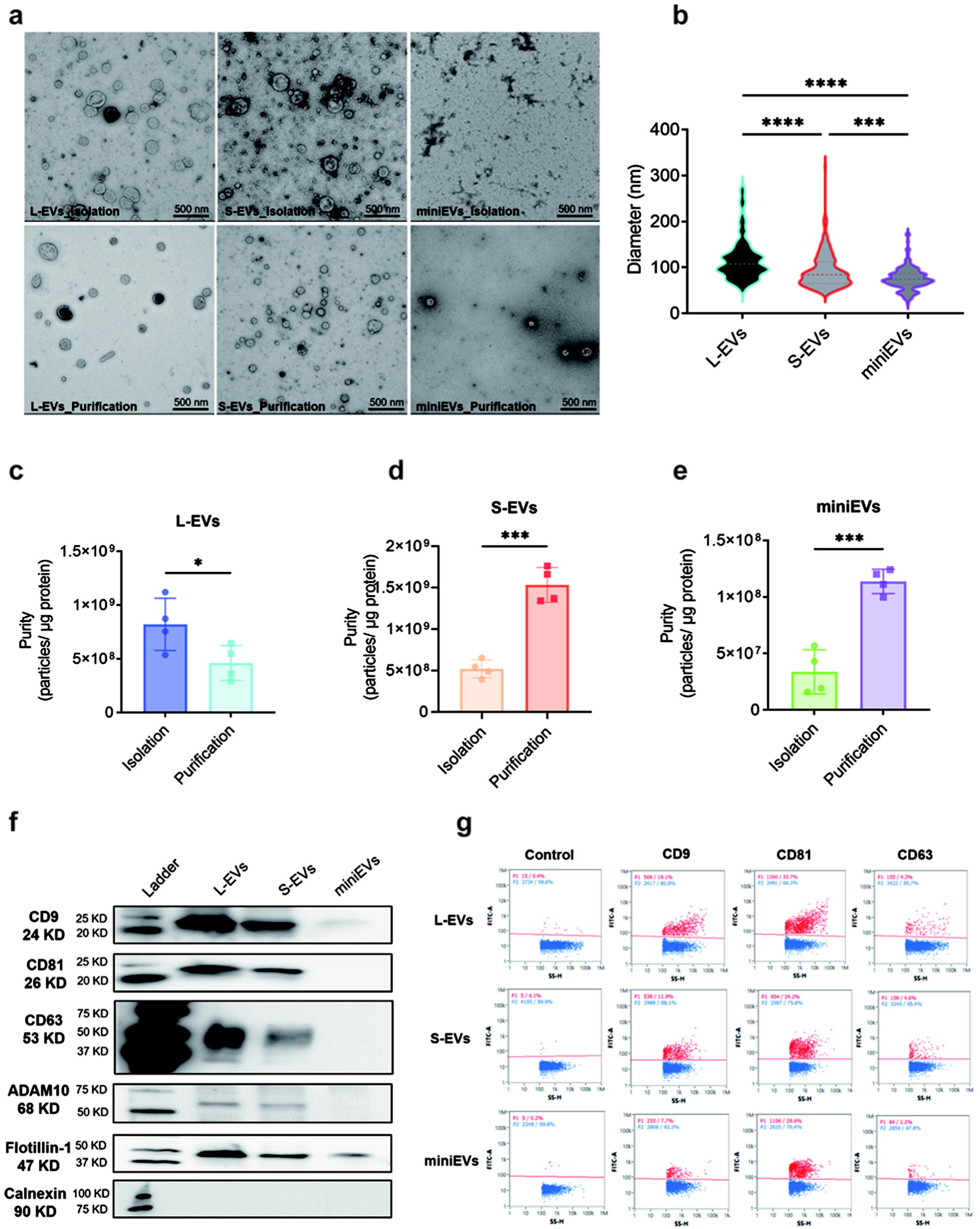
Characterization of three subpopulations of HEK293F-derived extracellular vesicles. **a)** The morphology of L-EVs, S-EVs, and miniEVs from HEK293F after ultracentrifugation (called “isolation” – upper panels) and after iodixanol density gradient (called “purification” – lower panels), visualized by negative-stained transmission electron microscopy (TEM). **b)** Diameter of HEK293F EVs measured manually on TEM images. **c-e)** The purity (measured as particles per protein) of EVs after different ultracentrifugation (called “isolation”) and after iodixanol density gradient (called “purification”). **f)** Western blot for CD9, CD81, CD63, ADAM10, Flotillin-1 and Calnexin in HEK293F EVs. **g)** Surface expression of CD9, CD81 and CD63 in HEK293F EVs measured by Nano-FCM. Data were analyzed using one-way ANOVA (Figure 1b) and student *t* test (Figure 1c-e). *, *P* <0.05; **, *P* < 0.01; ***, *P* < 0.001; ****, *P* < 0.0001. Abbreviations: EVs, extracellular vesicles; L-EVs, large EVs; S-EVs, small EVs.

Western blot analysis showed that the L-EVs and S-EVs carry comparable expression levels of CD9, CD81, CD63, ADAM10, and Flotillin-1 (Figure 1f). By contrast, the miniEVs showed lower expression of these markers per quantity of proteins. Notably, Calnexin, a marker of the endoplasmic reticulum, was undetectable in all three subpopulations (Figure 1f). Further, single-vesicle analysis by Nano-Flow cytometry (Nano-FCM) confirmed the presence of CD9, CD81, and CD63, but in slightly lower numbers in miniEVs vs L-EVs and S-EVs (Figure 1g). Together, these results indicate that three subpopulations of EVs can be isolated from HEK293F cells, which differ in size and protein cargos.

### HEK293F miniEVs, but not L- or S-EVs have anti-inflammatory properties

Next, we sought to evaluate the anti-inflammatory effect of the three subpopulations of EVs. Macrophages (RAW264.7) were exposed to bacterial outer membrane vesicles (OMV) to induce an innate immunity response, quantified by IL-6 and TNF-α release into the supernatant. Cells were then treated with EVs for 16 hours, followed by cytokine quantification in cell supernatant (Figure 2a). MiniEVs, but not the L-EVs or the S-EVs, showed significant reductions in IL-6 and TNF-α release (Figure 2b-c). Further, the miniEVs showed a dose-dependent effect on the cytokine release (Figure 2d-e). These data thus show that the three subpopulations have different biological activity, with the mini-EVs conveying anti-inflammatory function.

**Figure 2.**
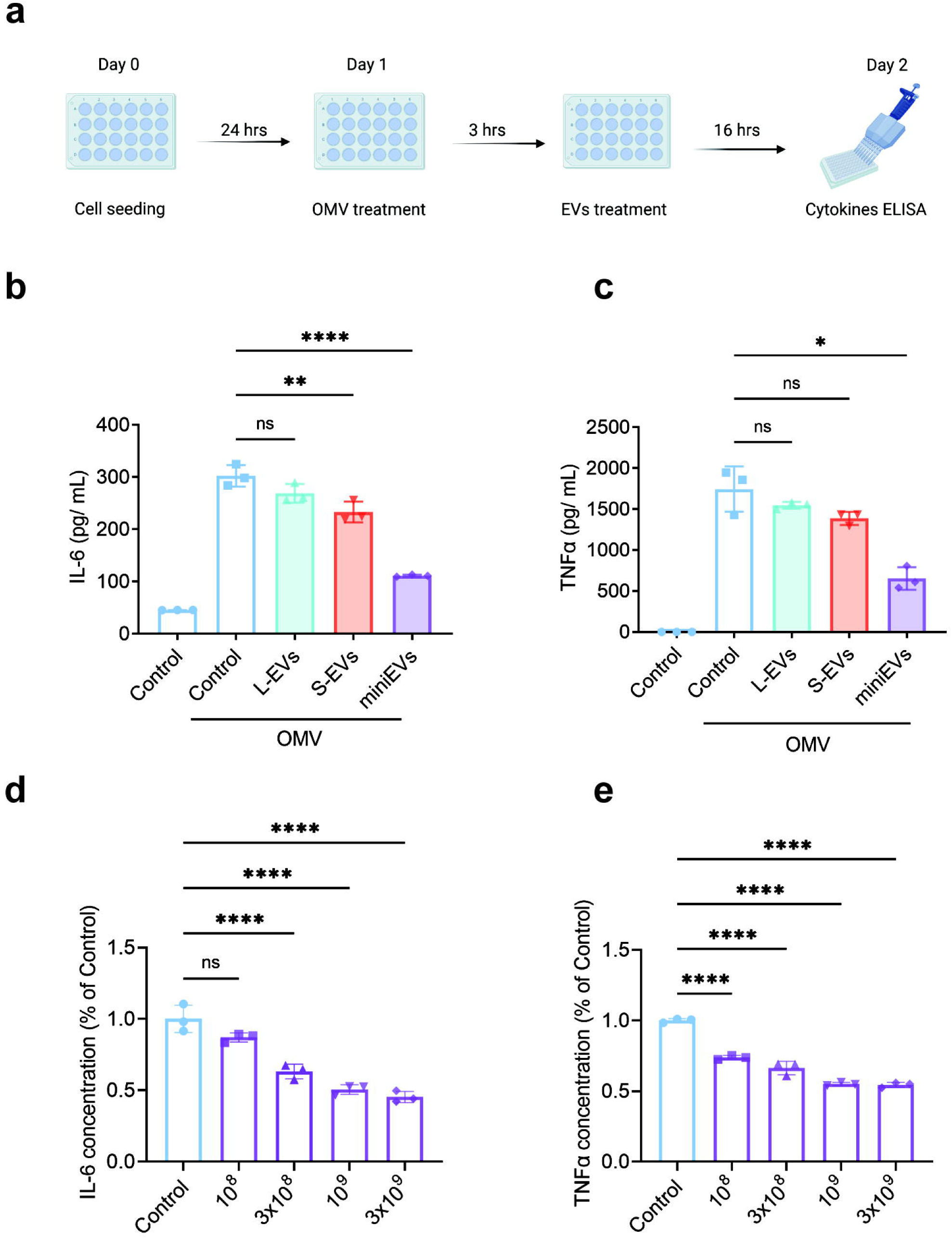
Anti-inflammatory properties of miniEVs from HEK293F cells. **a)** Experimental design: RAW 264.7 cells were exposed to E. coli outer membrane vesicles (OMV, 100 ng/mL) and subsequently treated with the different HEK293F EVs (10^9^/mL). **b-c)** The concentration of released IL-6 and TNF-α in the supernatant after exposure to OMV followed by HEK293F EVs treatment. **d-e)** Dose response inhibition by HEK293F miniEVs of IL-6 and TNF-α after exposure to OMV followed by HEK293F miniEVs treatment. Data were analyzed using one-way ANOVA. ns, no significant; *, *P* <0.05; **, *P* < 0.01; ****, *P* < 0.0001.

### Syndecan-4 is enriched in miniEVs

We conducted quantitative proteomic analysis (Tandem Mass Tag [TMT]) on the L-, S-, and miniEVs. The protein expression levels of the common EV proteins CD9, CD81, CD63, ADAM10, Flotillin-1, and Calnexin were assessed based on their abundance, showing higher amounts in L-EVs and S-EVs compared to miniEVs (Extended Data Figure 2), confirming Nano-FCM data (Figure 1g). A principal component analysis (PCA) revealed distinct protein expression profiles among the L-, S-, and miniEVs (Figure 3a). Specifically, 530 and 344 proteins were upregulated in mini-EVs compared to L-EVs and S-EVs, respectively (Figure 3b-c). A Venn diagram analysis identified 251 proteins that are upregulated in the miniEVs vs both L-EV and S-EV (Figure 3d). Of the 251 proteins, approximately 15% are integral membrane proteins, further arguing for the vesicular nature of the miniEVs. A Gene Ontology (GO) analysis for “biological process” of these 251 proteins revealed that five of the top ten pathways associated with these proteins were related to proteoglycans (Figure 3e). Specifically, we focused on proteoglycan proteins and could identify the presence of Syndecan-4 (SDC4) in the miniEVs, a membrane protein with potential anti-inflammatory effects (Figure 3f). Further, a sandwich ELISA for SDC4 showed that this molecule was present in significantly higher concentrations in miniEVs compared to L-EVs and S-EVs (Figure 3g). These data thus suggested that SDC4 potentially could contribute to the anti-inflammatory function of the miniEVs.

**Figure 3.**
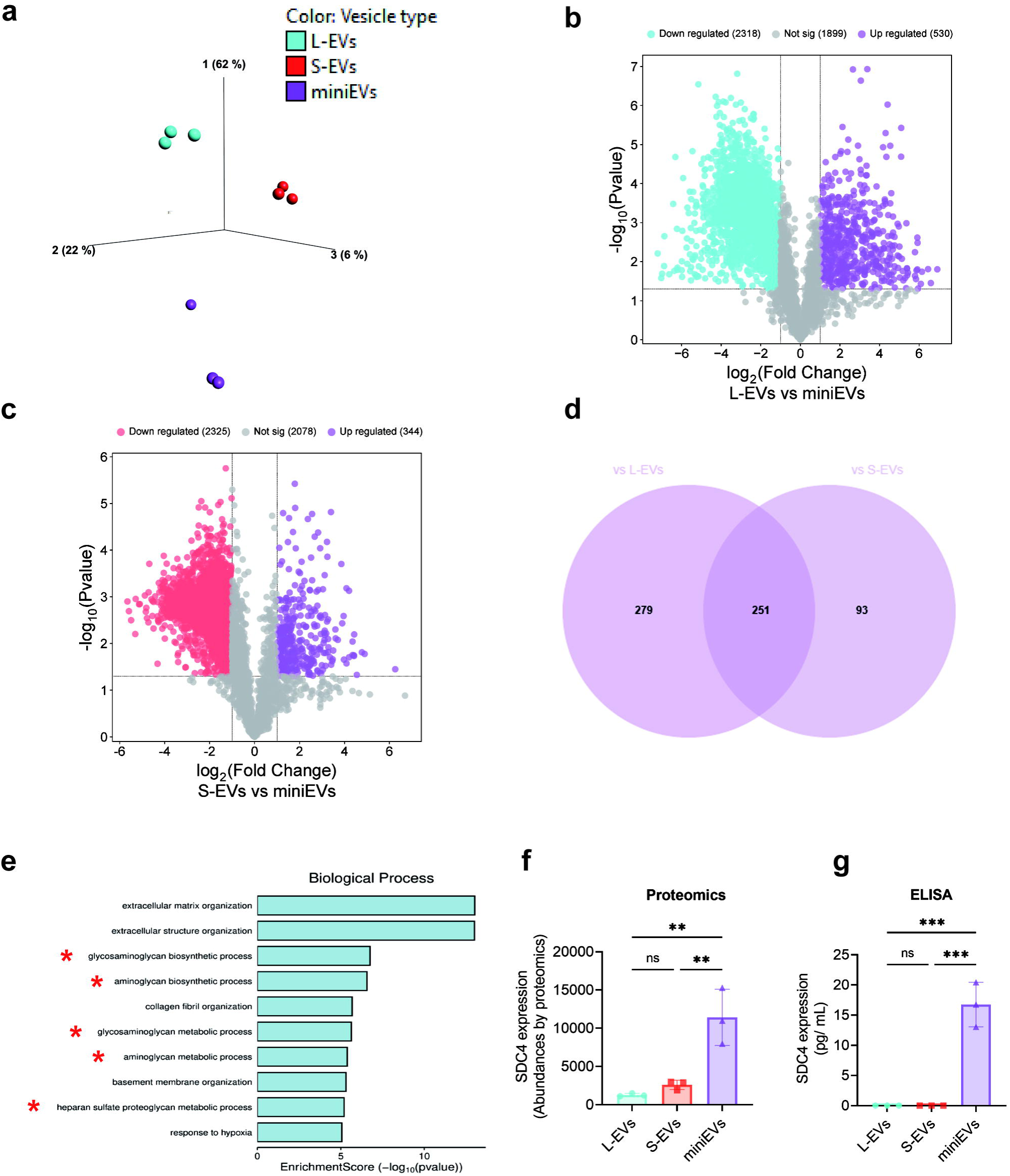
Proteomics analysis of subpopulations of EVs from HEK293F. Quantitative proteomics was used to determine the differences in the proteomes of the three subpopulations of EVs. **a)** Principal component analysis demonstrating the relationship of HEK293F-derived L-EVs, S-EVs, and miniEVs. **b-c)** Volcano plot illustrating differential expressed proteins between L-EVs and miniEVs (b) and between S-EVs and miniEVs (c). **d)** Venn diagram of the proteins that were shown to be upregulated in miniEVs vs L-EVs and in miniEVs vs S-EVs in Figure 3b and c. **e)** The most enriched Gene ontology terms (within the biological process category) associated with the proteins that were shown to be enriched in miniEVs in comparison with both L-EVs and S-EVs in Figure 3d. The ten most enriched terms (based on *p*-value) are displayed. f) Syndecan-4 (SDC4) abundance in the proteomics data. **g)** ELISA of SDC4 expression in 10^10^ HEK293F L-EVs, S-EVs and miniEVs. Data were analyzed using one-way ANOVA (Figure 3f-g). **, *P* < 0.01; ***, *P* < 0.001.

### Syndecan-4 overexpression induces anti-inflammatory properties in all HEK293F EVs

We hypothesized that an anti-inflammatory function could be induced in HEK293F L- or S-EVs by overexpression of SDC4. The processes of transfection and clone selection are detailed in Extended Data Figure 3a-d. Ultimately, one pure clone was chosen after a two-step selection process (Extended Figure 3d-e).

The expression levels of SDC4 were then compared in EVs isolated from the wild-type cells vs the SDC4-engineered cells, revealing a significant upregulation of SDC4 in all three subpopulations of HEK293F-SDC4 EVs as quantified by ELISA (Figure 4a), with the highest expression being observed in S-EVs. Furthermore, we confirmed that the L-EVs and S-EVs from the SDC4-expressing HEK293F cells, maintained morphology (Extended Figure 4a) and exhibited anti-inflammatory function (Extended Figure 4b-e). Therefore, from here on, we continued to isolate L- and S-EVs together from the HEK293-SDC4 cells, thus containing a mixture of L-EVs and S-EVs, as demonstrated by electron microscopy (Figure 4b). In these isolates, the SDC4 was highly overexpressed in SDC4 EVs vs wild-type HEK293F EVs, as shown by ELISA and Nano-FCM (Figure 4c-d). Furthermore, we confirmed the maintained anti-inflammatory function of the SDC4 EVs in macrophages. The SDC4 EVs significantly reduced IL-6 and TNF-α in the cell culture supernatant, compared to wild-type HEK293F EVs (Figure 4e-f). Further, the anti-inflammatory effect of the SDC4 EVs was found to be dose-dependent (Figure 4g-h). Together, these results confirm that HEK293F EVs can be engineered to become anti-inflammatory by overexpression of SDC4.

**Figure 4.**
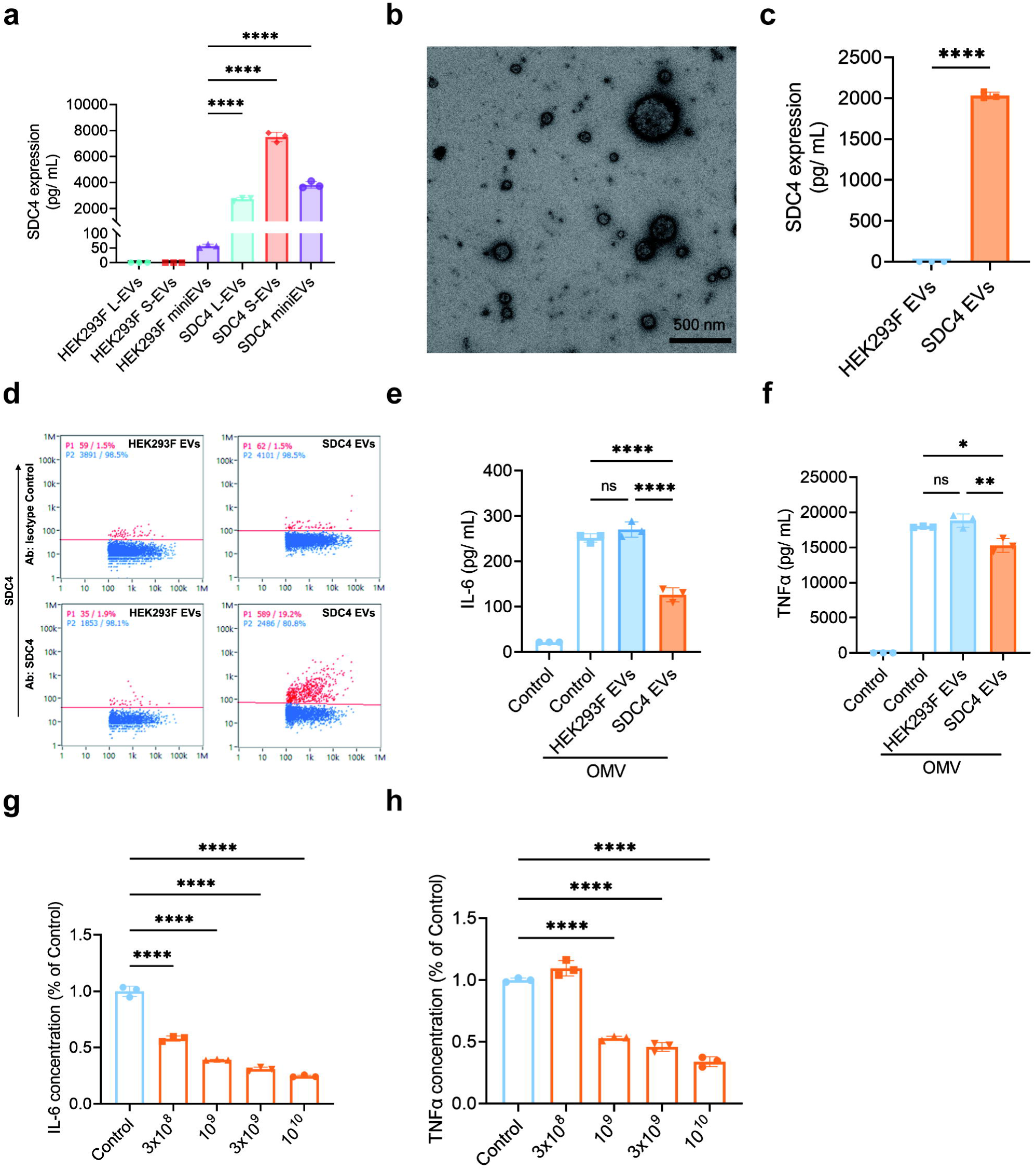
Characterization and anti-inflammation effects of SDC4 overexpressing HEK293F EVs. **a)** ELISA quantification of SDC4 expression in 10^10^ EVs from L-EVs, S-EVs and miniEVs isolated from wild-type or SDC4 overexpressing HEK293F cells. **b)** Representative TEM image of SDC4 overexpressing HEK293F EVs (SDC4 EVs). **c-d)** ELISA (c) and NanoFCM (d) analysis of SDC4 expression on HEK293F wild-type EVs and SDC4 EVs. **e-f)** The concentration of released IL-6 and TNF-α in the supernatant of RAW 264.7 cells after exposure to OMV (100 ng/mL) followed by treatment with HEK293F EVs (10^9^/mL) or SDC4 EVs (10^9^/mL). **g-h)** Dose-response curve for the inhibition of IL-6 (g) and TNF-α (h) release from RAW 264.7 cells after OMV treatment followed by increasing doses of SDC4 EVs. Data were analyzed using student *t* test (Figure 4c) and one-way ANOVA (Figure 4a, 4e-h). ns, no significant; *, *P* <0.05; **, *P* < 0.01; ****, *P* < 0.0001.

### Role of heparan sulfate for the anti-inflammatory effects of SDC4 EVs

Since SDC4 belongs to the heparan sulfate proteoglycans family (HSPGs) [22], we aimed to determine whether the overexpression of SDC4 changes the expression of heparan sulfate (HS). Firstly, overexpressing SDC4 on HEK293F cells increased their HS content (Extended Figure 5). Further, the EVs derived from the HEK293F-SDC4 cells also had increased HS content compared to wild-type EVs, as shown with both western blot and ELISA (Figure 5a-b). Next, we treated the SDC4 EVs with heparinase treatment, which maintained EV morphology (Figure 5c) but reduced EV HS levels according to western blot and ELISA (Figure 5d-e). Lastly, the anti-inflammatory effects of the SDC4 EVs was significantly reduced after heparinase treatment (Figure 5f-g). This suggests that the HS expression associated with the SDC4 overexpression mediates the anti-inflammatory effect of the engineered EVs.

**Figure 5.**
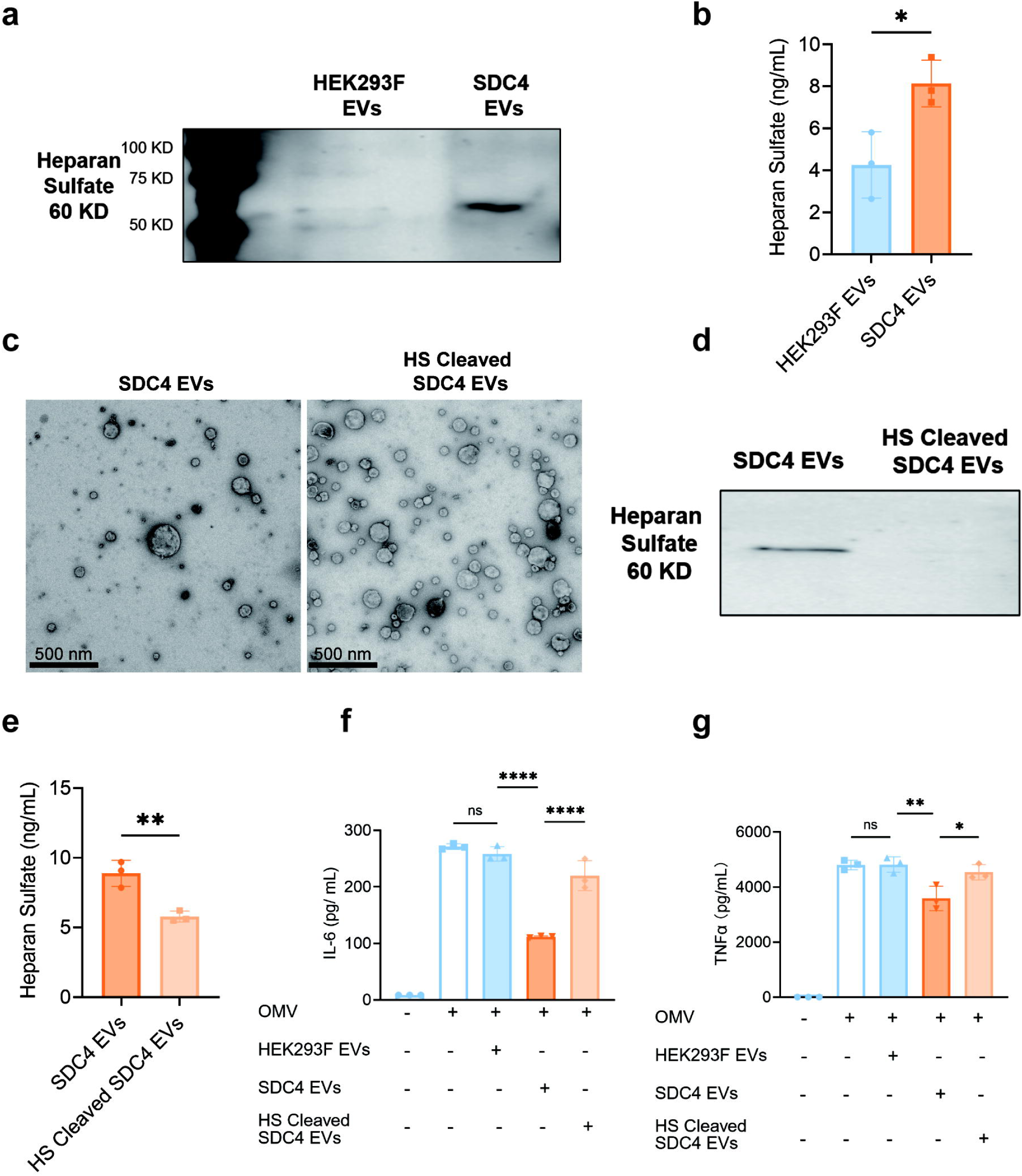
Role of heparan sulfate for the anti-inflammatory effects of SDC4 EVs. **a)** Western blot of heparan sulfate in wild-type HEK293F EVs and SDC4 EVs. 4 × 10^9^ EVs were used for the analysis. **b)** ELISA of heparan sulfate concentration in wild-type HEK293F EVs and SDC4 EVs. 10^10^ EVs were used for the analysis. **c)** Representative TEM images of SDC4 EVs untreated or treated with heparinase. **d)** Western blot of heparan sulfate in heparanase untreated or treated SDC4 EVs. 4 × 10^9^ EVs were used for the analysis. **e)** ELISA of heparan sulfate in heparanase-untreated or treated SDC4 EVs. 10^10^ EVs were used for the analysis. **f-g)** IL-6 and TNF-α concentration in the supernatant of OMV exposed RAW 264.7 cells treated with wild-type HEK293F EVs (10^9^/mL), or heparanase untreated or treated SDC4 EVs (10^9^/mL). Data were analyzed using student *t* test (Figure 5b, 5e) and one-way ANOVA (Figure 5f-g). ns, no significant; *, *P* <0.05; **, *P* < 0.01; ****, *P* < 0.0001.

### *In vivo* effects of SDC4 EVs in peritonitis

To determine whether the SDC4 EVs were also anti-inflammatory *in vivo*, we utilized a peritonitis model where mice are exposed to bacterial OMV by intraperitoneal (i.p.) injection (Figure 6a), and the mice were treated with either wild-type or SDC4 EVs locally. The SDC4 EV-treated group had a reduction in IL-6 expression in both the local peritoneal fluid and serum compared to mice treated with wild-type HEK293F EVs (Figure 6b-c).

**Figure 6.**
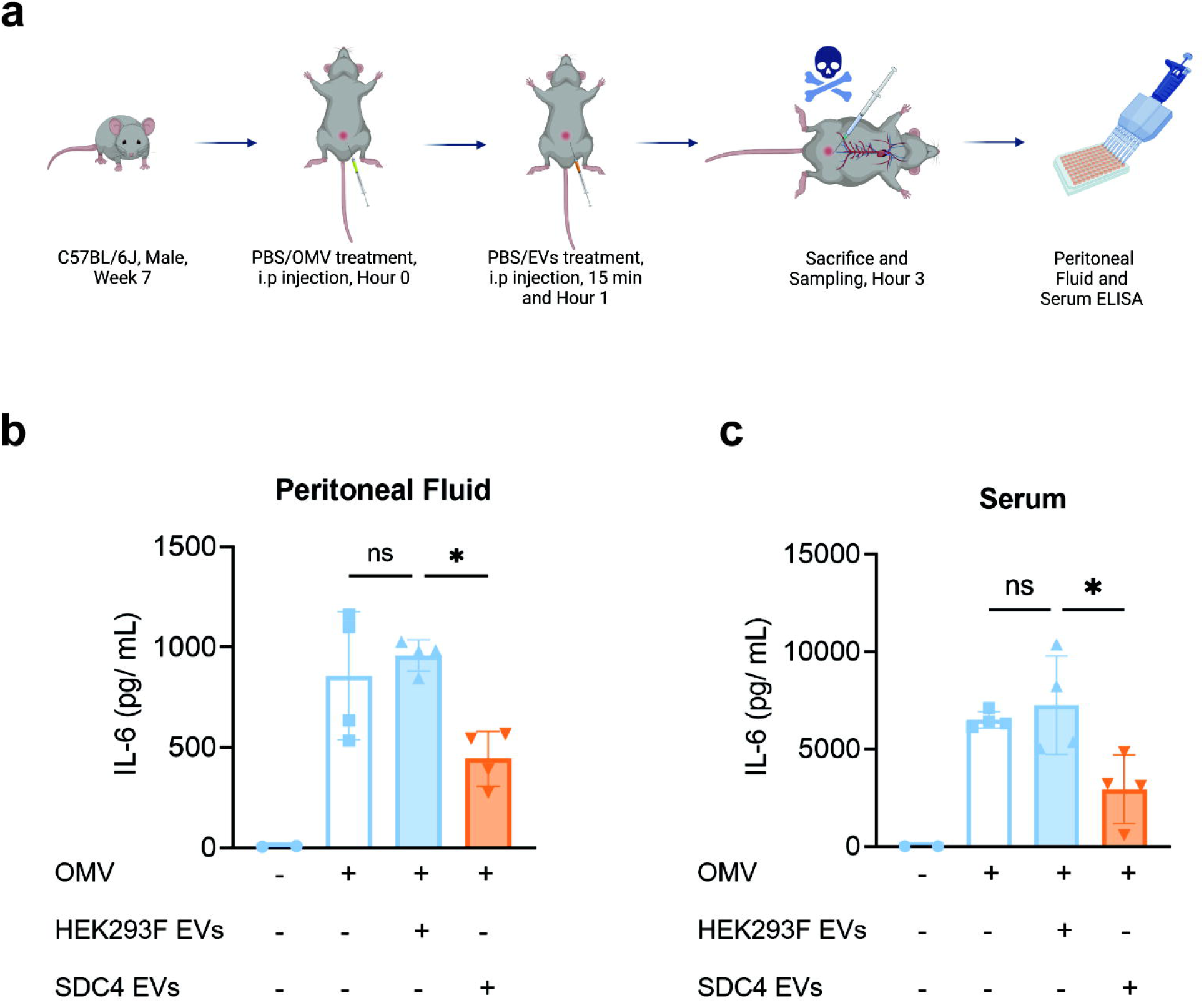
Effects of intraperitoneal (i.p.) administration of SDC4 EVs in mice exposed to i.p. OMV. **a)** Experimental design. Mice were treated OMVs (10 μg/mice) followed by treatment with wild-type HEK293F EVs (5×10^9^ EVs/mice/time point) and SDC4 EVs (5×10^9^ EVs/mice/time point) **b-c)** ELISA of IL-6 in mouse peritoneal fluid (b) and serum (c) after the OMV treatment followed by treatment with wild-type HEK293F EVs or SDC4 EVs. Data were analyzed using one-way ANOVA. ns, no significant; *, *P* <0.05; **, *P* < 0.01.

## Discussion

This study reveals that a seemingly distinct subpopulation of EVs released by HEK293F cells, which we have termed miniEVs, exhibits anti-inflammatory properties. This is in contrast to the larger EVs from HEK293F, lacking such functionality. Proteomic and bioinformatic analyses demonstrated that miniEVs possess a unique protein profile, including the presence of anti-inflammatory proteins absent in the larger EV subpopulations. Notably, we identified SDC4 in miniEVs, a membrane protein previously implied to regulate inflammation and belongs to HSPGs [22]. We therefore generated a stable HEK293F cell line overexpressing SDC4 to determine whether this molecule would be expressed in larger EVs, and whether it create a broader anti-inflammatory function of HEK293F EVs. Both miniEVs and larger EVs from a selected clone exhibited significantly elevated SDC4 levels and showed anti-inflammatory effects in a macrophage inflammation model. The SDC4-EVs had increased concentration of HS on their surface, and treatment with heparinase reduced the anti-inflammatory effects. Furthermore, in an *in vivo* peritonitis model, SDC4 EVs reduced inflammation. We conclude that HEK293F cells release a subpopulation of EVs (miniEVs) that are anti-inflammatory, and that this effect may be conveyed by SDC4. We confirmed that overexpression of SDC4 in EVs makes them anti-inflammatory.

Traditionally, EVs have been categorized based on their size and biogenesis into large extracellular vesicles (L-EVs) and small extracellular vesicles (S-EVs), commonly referred to as microvesicles and exosomes, respectively [21, 23–26]. Even though the EV subpopulations are released by the same cells, they may have fundamentally different functionalities [27], and specifically, different subpopulations of EVs from dendritic cell cultures have different effects on T-cells [28]. In this study, we explored the potential presence of EVs even smaller than the standard classifications by isolating EVs from the supernatant remaining after standard 2.5 hours of ultracentrifugation. We identified EVs with a smaller mean size than S-EVs, which we termed miniEVs, with round vesicle-structure as identified by electron microscopy, and they expressed EV membrane molecules such as tetraspanins. This makes the miniEVs distinct from non-EV structures such as Exomeres and Supermeres, which lack membrane structure [29–31]. Proteomic and principal component analyses further supported the uniqueness of miniEVs, with their molecular cargo rich in membrane proteins and suggesting anti-inflammatory functionality.

EVs derived from HEK293F cells have been extensively studied for their potential as drug delivery vehicles due to their biocompatibility and ability to target specific cells [32–35]. In this study, we showed miniEVs exhibited ∼60% anti-inflammation efficacy at the 10^9/mL concentration, which is the similar efficacy with MSC-EVs [36]. Our previous study has documented that the membrane components of MSC-EVs play a vital role in their anti-inflammatory effects [36], and we therefore hypothesized that miniEVs also may mediate their effects via membrane components. Indeed, proteomic profiling revealed distinct membrane protein cargo in miniEVs, with enrichment of proteins involved in extracellular matrix organization, glycan biosynthesis, and HS processes. Notably, SDC4 was significantly upregulated in miniEVs, with low expression in L-EVs and S-EVs.

SDC4 is known to regulate inflammation, with evidence supporting both pro- and anti-inflammatory functions depending on the biological context. For instance, inhibition of SDC4 signaling alleviates asthma features by modulating immune cell migration and activation, highlighting a potential pro-inflammatory role [37]. Similarly, in autoimmune arthritis, SDC4 regulates B cell migration and germinal center formation, and deletion of SDC4 reduces B cell migration and disrupts germinal center formation. This reduction of SDC4 ultimately resulted in attenuation of the arthritis severity in this experimental model [38]. In contrast, SDC4 can also exhibit potent anti-inflammatory properties in other disease models. For example, SDC4-deficient mice show increased mortality following lipopolysaccharide (LPS) challenge compared to wild-type controls, suggesting a protective, anti-inflammatory role in this context [39]. Similarly, in LPS-induced lung injury, SDC4 deficiency exacerbates neutrophilic inflammation and CXC chemokine release, further supporting an anti-inflammatory function [40]. SDC4 deficiency also promotes macrophage-driven inflammation in atherosclerosis, suggesting its potential as a therapeutic target for disease prevention [41]. These, to some degree, contradictory findings underscore a potential dual role of SDC4 in orchestrating inflammatory responses. Importantly, we have identified the anti-inflammatory effects of treatment with SDC4-expressing EVs in innate immunity responses *in vitro* and *in vivo*, and focused on testing its potential therapeutic efficacy *in vitro* and *in vivo*.

Based on these considerations, we have overexpressed SDC4 in HEK293F cells, resulting in SDC4 expression in all EV subpopulations, which in turn conveys anti-inflammatory properties to the L-EVs and S-EVs. SDC4 expression in EVs also resulted in increased HS levels on their surface. We therefore hypothesized that this glycosaminoglycan could mediate the SDC4 effects. Indeed, the enzymatic reduction of HS significantly reduced the anti-inflammatory efficacy of the SDC4 EVs. While residual anti-inflammatory activity persisted, these findings highlight that HS can contribute to the observed anti-inflammatory effects of SDC4 overexpression on EVs. The anti-inflammatory effects of SDC4 EVs were confirmed *in vivo* in a murine peritonitis model, where treatment resulted in ∼50% reduction in inflammation, suggesting they could potentially be suitable to treat human inflammatory diseases.

SDC4 overexpression may, via HS, interact with a diverse array of ligands, including growth factors, adhesion molecules, cytokines, chemokines, proteinases, and other extracellular matrix proteins [42]. Further, HS is known to have both anticoagulant and anti-inflammatory properties [43, 44]. In this study, we extended these results by showing that SDC4 expression on EVs induces an anti-inflammatory functionality via HS. It is unclear whether the SDC4 EV-associated HS directly binds cytokines, acting as a molecular sponge [45], or whether other anti-inflammatory mechanisms are involved. It has been implicated that SDC4 also may regulate inflammatory signaling pathways via its cytoplasmic domain [46]. HS is known for its anti-inflammatory properties, but its therapeutic application is limited by challenges in its synthesis. Here, we suggest that overexpression of SDC4 on EVs may offer a viable strategy to harness the benefits of HS.

There are several limitations in this study that should be considered. Firstly, only 20% of the released EVs were found to express SDC4, and it is possible that achieving expression in a larger proportion of EVs could result in further enhanced anti-inflammatory effects. Further, more detailed evaluation of the potential pro-inflammatory vs the anti-inflammatory properties of SDC4 on EVs requires extra attention, before these EVs can be tested clinically.

In conclusion, this study identifies miniEVs as a novel EV subpopulation that carries SDC4, exhibiting unique anti-inflammatory properties. Engineering HEK293F cells to overexpress SDC4 yielded EVs with potent anti-inflammatory effects, which is associated with HS surface expression. These findings highlight the therapeutic potential of SDC4 EVs and underscore the importance of EV membrane components in modulating inflammation. These results underline the opportunity to engineer EVs to become anti-inflammatory, thereby creating a platform for future anti-inflammatory therapeutics.

## Online Methods

### Cell culture

We utilized two cell lines: HEK293F (Thermo Fisher Scientific) and RAW 264.7 (ATCC). HEK293F cells were cultured in Freestyle 293 medium (Thermo Fisher Scientific), either in suspension for EVs production or adherently for plasmid transfection and clone selection. Suspension cultures were incubated at 37°C with 70% humidity and 5% CO2, shaking at 130 rpm, while adherent cultures included 10% FBS. RAW 264.7 cells were maintained in Dulbecco’s modified Eagle’s medium (Cytiva Hyclone) with 10% FBS, 100 U/mL penicillin, and 100 µg/mL streptomycin. All cultures were kept at 37°C with 5% CO2.

### Extracellular vesicles (EVs) isolation and purification

A combined ultracentrifugation and iodixanol gradient cushion methods were applied for EVs isolation (Extended Data Figure 1). Cell culture media was centrifuged at 300 × g for 10 minutes and 2000 × g for 20 minutes to remove cells, debris, and large apoptotic bodies. The supernatant was ultracentrifuged at 14,500 × g for 20 minutes for isolation of L-EVs (16,500 rpm, Type 45 Ti fixed angle rotor, 1278 as k-factor, Beckman Coulter), 118,000 × g for 2.5 hours for isolation of S-EVs (38,500 rpm, Type 45 Ti fixed angle rotor, 181 as k-factor, Beckman Coulter), and 118,000 × g for 16 hours for isolation of miniEVs (38,500 rpm, Type 45 Ti fixed angle rotor, 181 as k-factor, Beckman Coulter). The pellet was resuspended in PBS and further purified with an iodixanol density cushion, followed by ultracentrifugation at 100,000 × g for 2 hours(28,000 rpm, SW 41 Ti swinging rotor, 265 as k-factor, Beckman Coulter). The EVs were collected at the 10-30% iodixanol interphase.

### Transmission Electron Microscopy (TEM)

TEM was employed to identify and characterize EVs in this study and non-fixed negative staining was performed prior to TEM. For negative staining, EVs were applied to g low-discharged 300-mesh copper grids (Electron Microscopy Sciences) for 30 seconds, followed by two washes with water. Subsequently, the EVs were stained with 2% uranyl formate for 1 minute. The negatively stained EVs were then examined using a TALOS L 120C transmission electron microscope (Thermo Fisher Scientific) operating at 120 kV, equipped with a BM-Ceta CMOS 4k × 4k CCD camera.

### Nanoparticle Tracking Analysis (NTA)

The EV particle concentration was measured using a ZetaView analyzer (Particle Metrix) with eleven data points. Camera sensitivity was set to 80 for Large/Small EVs and 85 for miniEVs. Data interpretation used ZetaView software (version 8.05.12 SP1) with brightness settings from 30 to 1000.

### Western Blot

Protein concentrations of cell lysates and EVs were measured using a Pierce BCA Protein assay kit (Thermo Fisher Scientific). EVs (5 μg or 4 × 10^9^ particles) or cell-extracted proteins (10 μg) were diluted 1:3 in loading buffer (Bio-Rad), heated at 95°C for 5 minutes, and loaded onto 10-20% gels (Bio-Rad) for electrophoresis. The gels were run at 80 V for 20 minutes and 180 V for 40 minutes, then transferred to membranes with a semi-dry transfer chamber (Bio-Rad). Membranes were blocked for 60 minutes, incubated with primary antibodies overnight at 4°C, washed, incubated with secondary antibodies for 60 minutes at room temperature, washed again, and exposed to a detection reagent for visualization. The primary antibodies were as follows: CD9 (EMD Millipore, 1:1000), CD81 (Abcam, 1:1000), CD63 (BD Pharmingen, 1:1000), ADAM10 (Rnd Systems, 1:500), Flotilin-1 (Abcam, 1:1000), Calnexin (Cell Signalling Technology, 1:1000), Heparan Sulfate (Sigma Aldrich, 1:1000), β-actin (Cell Signalling Technology, 1:1000).

### Nano-FCM

EV surface molecule expression was evaluated using a Flow NanoAnalyzer (NanoFCM Inc.) following the manufacturer’s instructions. Before loading samples, the analyzer was calibrated with QC beads, size beads, and a blank control. Events per minute were kept below 12,000; samples exceeding 2,000 events in the first 15 seconds were diluted to ensure counts between 11,000 and 12,000. For immunofluorescent staining, 1 µL of diluted antibodies was added to 4 µL of the diluted sample, incubated for 40 minutes at room temperature in the dark, then diluted 10-fold with DPBS before data acquisition. The antibodies used for the staining step were: CD9 (BD Pharmingen, FITC), CD81 (BD Pharmingen, FITC), CD63 (BD Pharmingen, FITC), Syndecan 4 (Human Syndecan-4 APC-conjugated Antibody, biotechne), Rat IgG2A Allophycocyanin Isotype Control (biotechne). Anti-CD9, anti-CD81 and anti-Syndecan 4 were diluted 6-fold, while anti-CD63 was diluted 2-fold.

### Proteomics

Each EV proteomic analysis sample was normalized to contain 40 µg of protein (9 samples total). Sodium dodecyl sulfate (SDS) was added to achieve a 2% concentration. Samples underwent a modified filter-aided preparation [47]: reduced with 100 mM DTT at 60°C for 30 minutes, filtered with Microcon-30kDa units, and washed with 8 M urea and digestion buffer (DB; 50 mM TEAB and 0.5% SDC). They were then alkylated with 10 mM methyl methanethiosulfonate in DB for 30 minutes at room temperature. Trypsin digestion occurred at a 1:100 ratio overnight at 37°C, with an additional trypsin portion added for 2 more hours. Peptides were collected, labeled with TMT 18-plex reagents (Thermo Fisher Scientific), combined into one set, and SDC was removed by acidification with 10% TFA. The TMT-set was further purified with a High Protein and Peptide Recovery Detergent Removal Spin Column, followed by Pierce Bioinformatic analysis of proteomics data.

### Generation of the stable Syndecan-4 (SDC4) clones, transient transfected clones and a pure clone

The stable SDC4 clone was created by transfecting human SDC4 membrane plasmid (Genescript) using the Lipofectamine 2000 system (Thermo Fisher Scientific), followed by two sequencing selections. Two days before transfection, 1.0 × 10^6^ cells were seeded in a 6-well plate with growth medium. On transfection day, the medium was refreshed, and a mixture of 4 µg plasmid and 10 µL Lipofectamine 2000 in 500 µL IMDM (Cytiva Hyclone) was incubated for 20 minutes, then added to the cells. A mock transfection was performed using the same mixture without the plasmid.

Antibiotic G418 (500 µg/mL) was added 48 hours post-transfection for selection. The growth medium was refreshed every 48 hours, with clone pools expanded or split as needed. A selection procedure was initiated when mock culture viability decreased, involving both the transfected and mock groups. Transfected cells were seeded in 10× dilutions on Petri dishes. After two weeks, 24 clones were picked and transferred to a 24-well plate with 500 µL of media per well. Clones were expanded and analyzed via flow cytometry for optimal membrane protein expression. The cells before the Petri dish selection were partly saved as transient transfection clones. The clone with the highest SDC4 expression and good proliferation was selected as the pure clone.

### Flow cytometry

To assess transfection efficiency and membrane protein expression, cells were analyzed using a BD FACS Verse flow cytometer with BD FACS Suite software (BD Biosciences). 100,000 cells were suspended in 1.5 mL ice-cold FACS buffer (1% FBS in PBS), pelleted, and resuspended in human IgG (1 mg/mL in D-PBS). After a 15-minute incubation, Human Syndecan-4 APC-conjugated Antibody (R&D Systems) and Rat IgG2A Isotype Control (R&D Systems) were added for 30 minutes. Cells were washed, resuspended in FACS buffer, and 10,000 events per sample were collected and analyzed using FlowJo software (Tree Star Inc, Ashland, OR, USA).

### Syndecan-4 (SDC4) ELISA

SDC4 expression on EV surfaces was assessed with a DuoSet ELISA kit (R&D Systems). Briefly, 10^9^ or 10^10^ particles were captured on a capture antibody-coated plate for 2 hours at room temperature, followed by a 2-hour incubation with a detection antibody. HRP and substrate were then added, and luminescence was measured with a plate reader. SDC4 expression was determined using a standard curve.

### Heparan Sulfate (HS) ELISA

Heparan sulfate was detected using a Sandwich-ELISA kit (Ambio). Standards or particles (10^9^ or 10^10^) were added to antibody-coated wells for 2 hours at 37℃. Biotinylated detection antibodies and Avidin-HRP were then added and incubated. After washing away free components, a substrate was added, turning blue in wells with Human HS, detection antibody, and Avidin-HRP. The reaction was stopped, turning the solution yellow, and the optical density (OD) was measured at 450 nm. Human HS concentration was determined by comparing sample OD to a standard curve.

### HS Cleaved SDC4 EVs

SDC4 EVs (8.35 × 10^10^), 1 U heparanase (Heparinase I and III Blend, Sigma-Aldrich), and PBS were combined into 1 mL and incubated at 30°C for 4 hours. Post-incubation, the EVs were washed via ultracentrifugation at 118,000 × g for 58 minutes at 4°C (53,000 rpm, TLA 100.3 fixed angle rotor, 51 as k-factor, Beckman Coulter) using a bench-top ultracentrifuge (Beckman Coulter).

### Isolation of outer membrane vesicles (OMV) derived from Escherichia coli

The bacterial cultures were centrifuged at 6,000 × g at 4°C for 20 minutes. The supernatant was filtered through a 0.45 μm vacuum filter and concentrated using a Vivaflow 200 ultrafiltration module (Sartorius) with a 100 kDa membrane. The concentrate was then ultracentrifuged at 150,000 × g at 4°C for 3 hours (Type 45 Ti fixed angle rotor, 181 as k-factor, Beckman Coulter) and resuspended in PBS.

### *In vitro* evaluation of the anti-inflammatory potentials of EVs

RAW 264.7 cells (10^5^) were seeded in a 24-well plate, treated with 100 ng/mL OMV for 3 hours, and then given EVs treatment for 16 hours. The supernatant was collected and centrifuged at 600 × g for 10 minutes and 3000 × g for 20 minutes at 4°C. IL-6 and TNF-α concentrations were measured using a DuoSet ELISA kit (R&D Systems).

### Animal studies ethics

C57BL/6J mice (6 weeks old) were obtained from Charles River and raised in the experimental animal room at the Experimental Biomedicine facility at the University of Gothenburg, Sweden. The local Animal Ethics Committee in Gothenburg approved the experiment (permit no. Dnr 5.8.18–10595/2023), conducted under institutional animal care guidelines.

### *In vivo* evaluation of the anti-inflammatory potentials of EVs

C57BL/6J mice (7 weeks old) received a 10 μg intraperitoneal dose of OMV to trigger peritonitis. HEK293F EVs and SDC4 EVs (5 × 10^9^) were administered via intraperitoneal injection at 15 minutes and 1 hour post-OMV administration. At 3 hours after OMV administration, mice were euthanized using xylazine chloride (10 mg/kg; Bayer) and ketamine hydrochloride (100 mg/kg; Pfizer AB). Blood and peritoneal fluid were collected, centrifuged twice at 2000 × g for 10 minutes at 4 °C. IL-6 levels were measured using a DuoSet ELISA kit (R&D Systems), with serum diluted 1:10 and peritoneal fluid undiluted.

### Statistical analysis

All data are presented as means ± standard deviation. Comparisons were performed using student *t* test and one-way ANOVA, as appropriate, using GraphPad Prism version 9.0. *P* < 0.05 was considered statistically significant (*, *P* < 0.05; **, *P* < 0.01; ***, *P* < 0.001; ****, *P* < 0.0001).

## Acknowledgements

This work was supported by the Swedish Research Council (Grant No. 2021-03538) and the Swedish Heart-Lung Foundation (Grant No. 20210707) to J.L.. This work was also supported by the International Cultivation Program for Excellent Young Talents of Guangdong Province and National Natural Science Foundation of China (Grant No. 82203711) to L.Y.. We acknowledge the Centre for Cellular Imaging at the University of Gothenburg and the National Microscopy Infrastructure (VR-RFI 2019-00217) for providing assistance in transmission electronic microscopy. The quantitative analysis was performed at the Proteomics Core Facility, Sahlgrenska Academy, Gothenburg University, with financial support from SciLifeLab and BioMS. Illustrations (Graphical abstract, Figure 2a, Figure 6a, Extended Data Figure 1a, Extended Data Figure 3a) were created with https://BioRender.com. We acknowledge Prof. Lei Zheng at Southern Medical University (China) and Prof. Roger Olofsson Bagge at Gothenburg University (Sweden) for their support for this work. We acknowledge Xiaoyu Wang at International School of Gothenburg Region for his useful discussion on the mini bubbles.

## Author contributions

LY, JL: Conceptualization. LY: Discovery of HEK293 miniEV functionality. LY, JL: designed the overall research approach. LY: performed the in vitro experiments. LY, MB: performed the cloning. LY, KSP: performed the in vivo experiments. LY, MB: performed the nano-FCM experiments. LY, CL: analyzed the proteomic data. LY, JL: analyzed and interpreted the data. MB, KSP, CL: provided input on experimental plan and discussion of results. LY, CL, JL: wrote the manuscript. MB, KSP: review and editing of the manuscript. JL: Supervised and led the study. JL, LY: acquired funding for the study (experiments and post-doc salary, respectively).

## Supplementary Information

Extended Figure 1. Method for enrichment of L-EVs, S-EVs, and miniEVs from conditioned medium from HEK293F cells.

Extended Figure 2. Protein expression of EV proteins in the proteomic analysis of the HEK293F EVs.

Extended Figure 3. The process of establishing an SDC4 expressing HEK293F clone.

Extended Figure 4. Characterization and anti-inflammation function of L-EVs, S-EVs and miniEVs of SDC4 overexpressing HEK293F cells.

Extended Figure 5 Western blot of heparan sulfate expression of wild-type HEK293F cells vs SDC4 overexpressing HEK293F cells.

