## Supplementary Information for "Development of Anti-Inflammatory Extracellular Vesicles by Surface Expression of Syndecan-4"

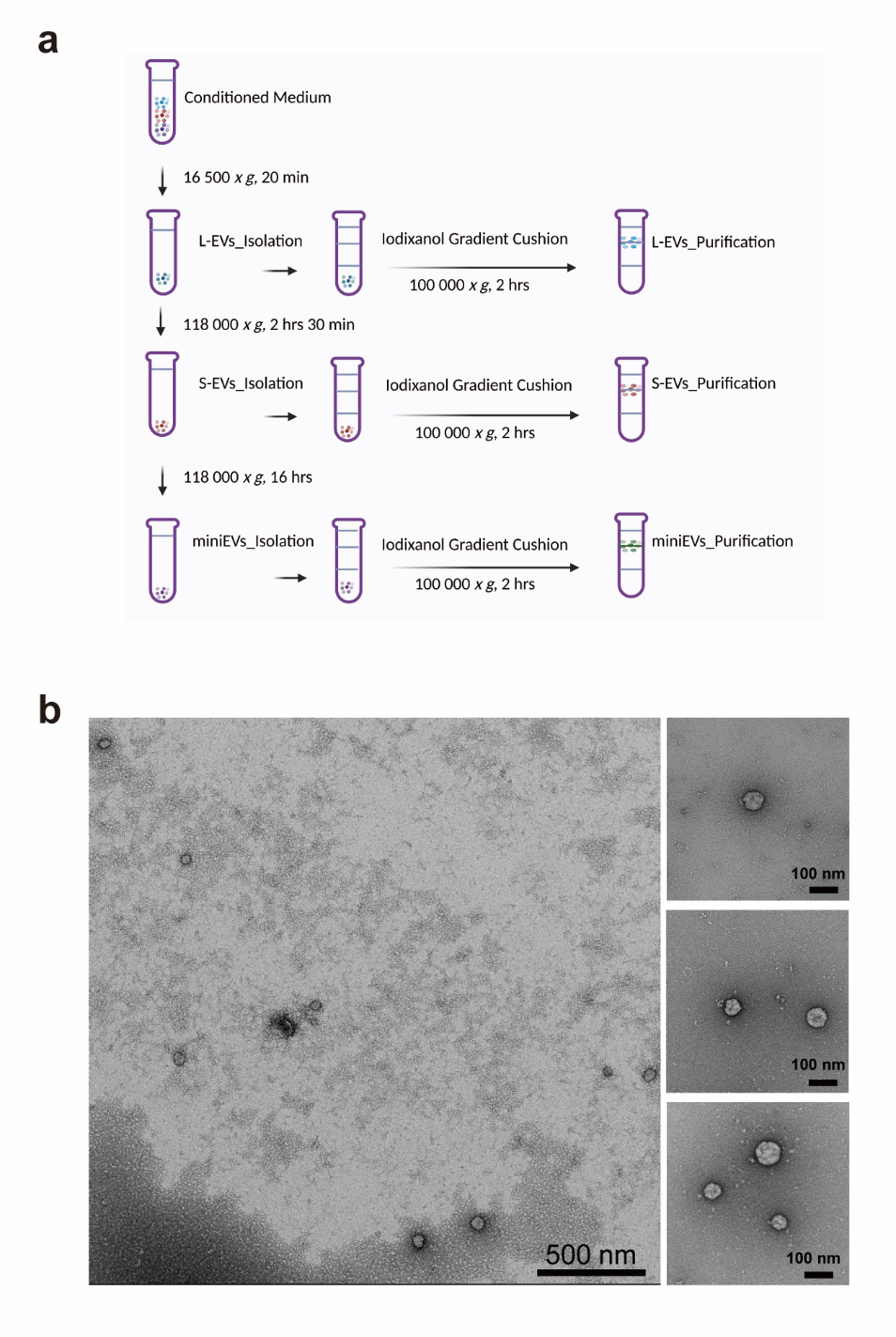

**Extended Figure 1. Method for enrichment of L-EVs, S-EVs, and miniEVs from conditioned medium from HEK293F cells.** **a)** The centrifugation methods for the enrichment of the different EVs. **b)** TEM images of HEK293F miniEVs at different magnification. Bigger image: 22,000 X magnification; Smaller images: 73, 000 X magnification.

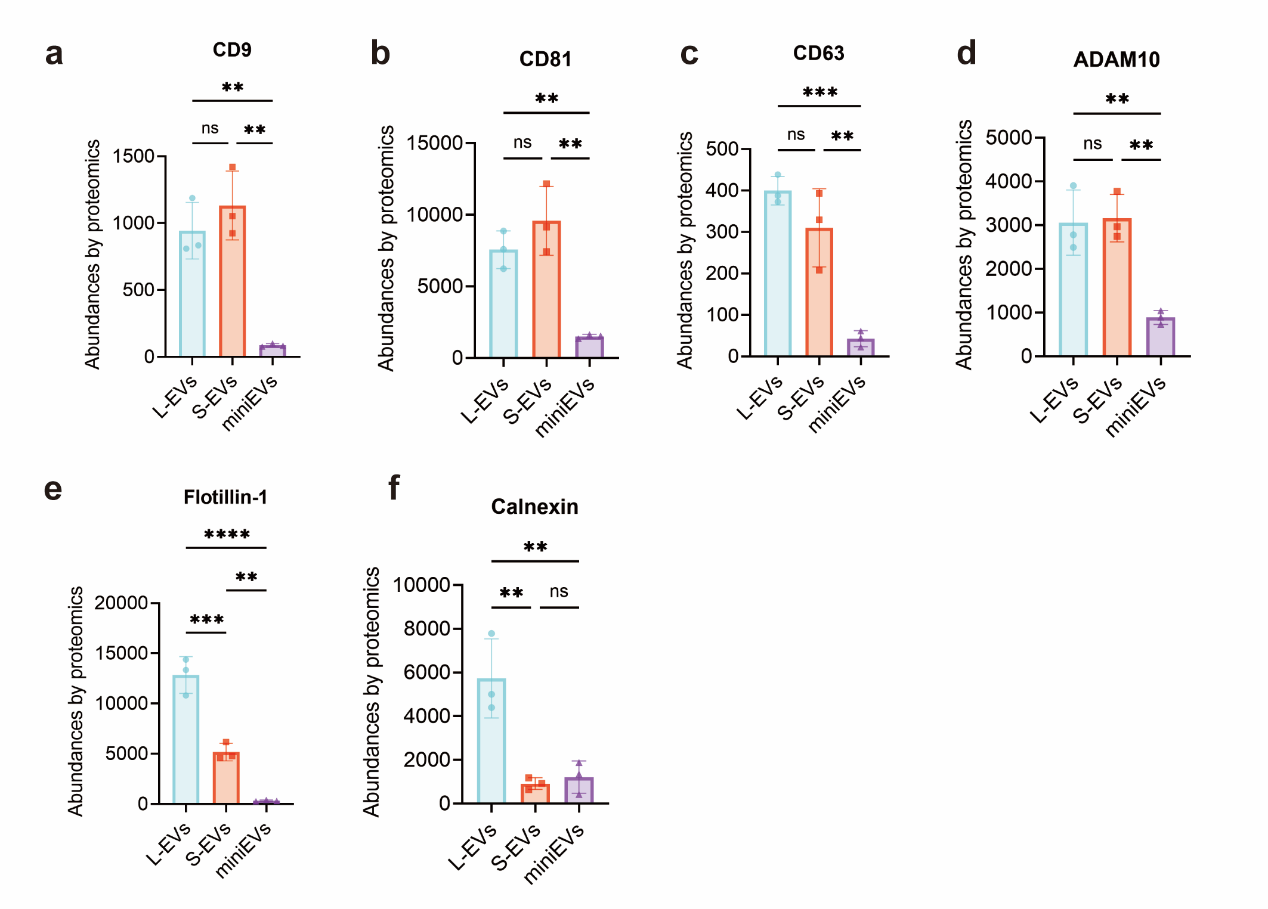

**Extended Figure 2. Protein expression of EV proteins in the proteomic analysis of the HEK293F EVs.** **a-e)** The abundance in the LC-MS/MS analysis for the commonly analyzed EV proteins; CD9 (a), CD81 (b), CD63 (c), ADAM10 (d), and Flotillin-1 (e). **f)** The abundance in the LC-MS/MS analysis for the endoplasmic reticulum protein Calnexin. Data were analyzed using one-way ANOVA. **, *P* < 0.01; ***, *P* < 0.001; ****, *P* < 0.0001.

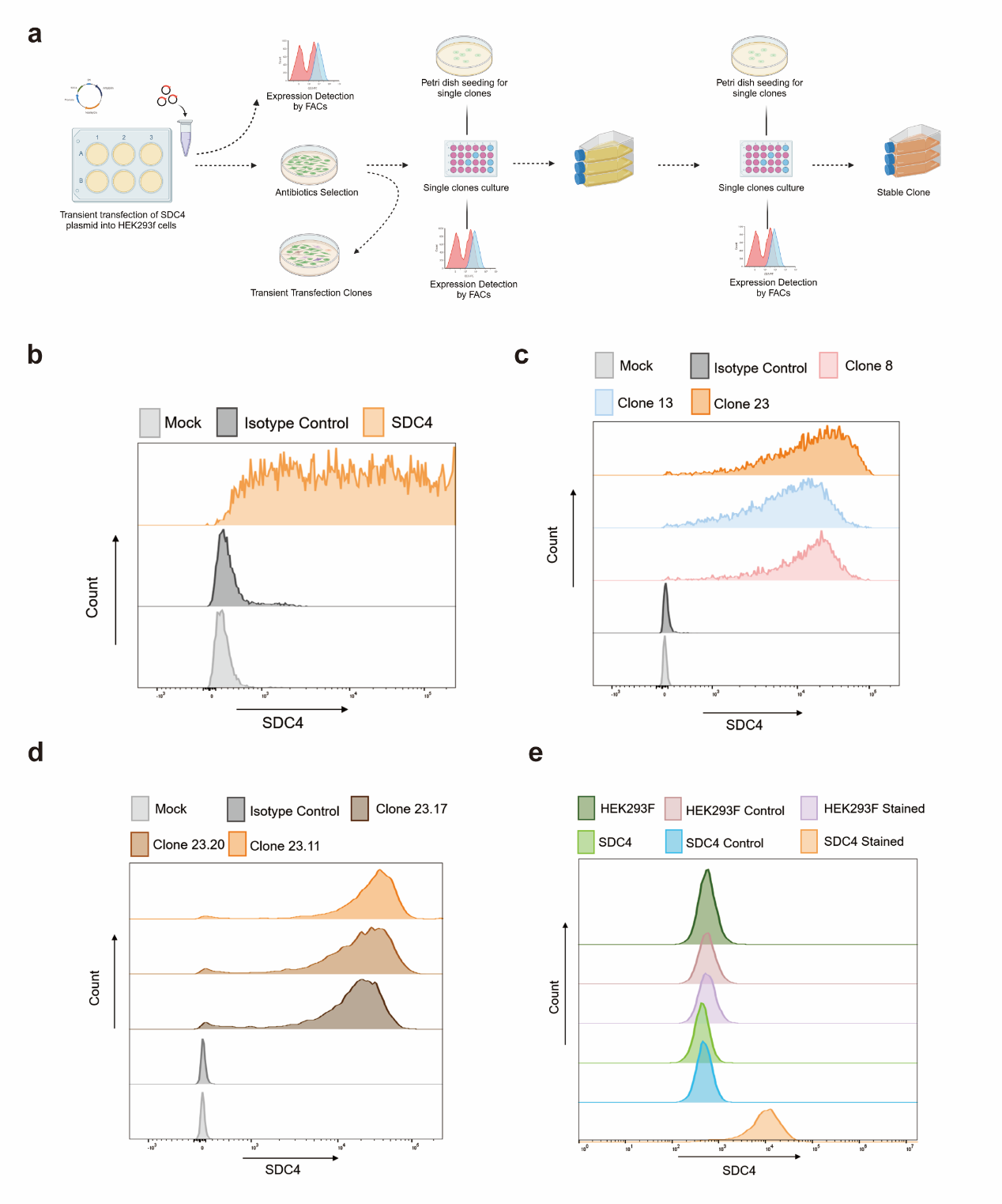

**Extended Figure 3. The process of establishing an SDC4 expressing HEK293F clone.** **a)** Experimental design for the transfection, which included the production of transient transfection clones and the selection of stable clones. **b)** Flow cytometry data of SDC4 expression on the transient transfection clones. **c)** Flow cytometry data of three clones after the first selection process. **d)** Flow cytometry data of the top three clones after the second selection process. **e)** Flow cytometry data comparing SDC4 expression of HEK293F wild-type cells and final selected SDC4 HEK293F clone.

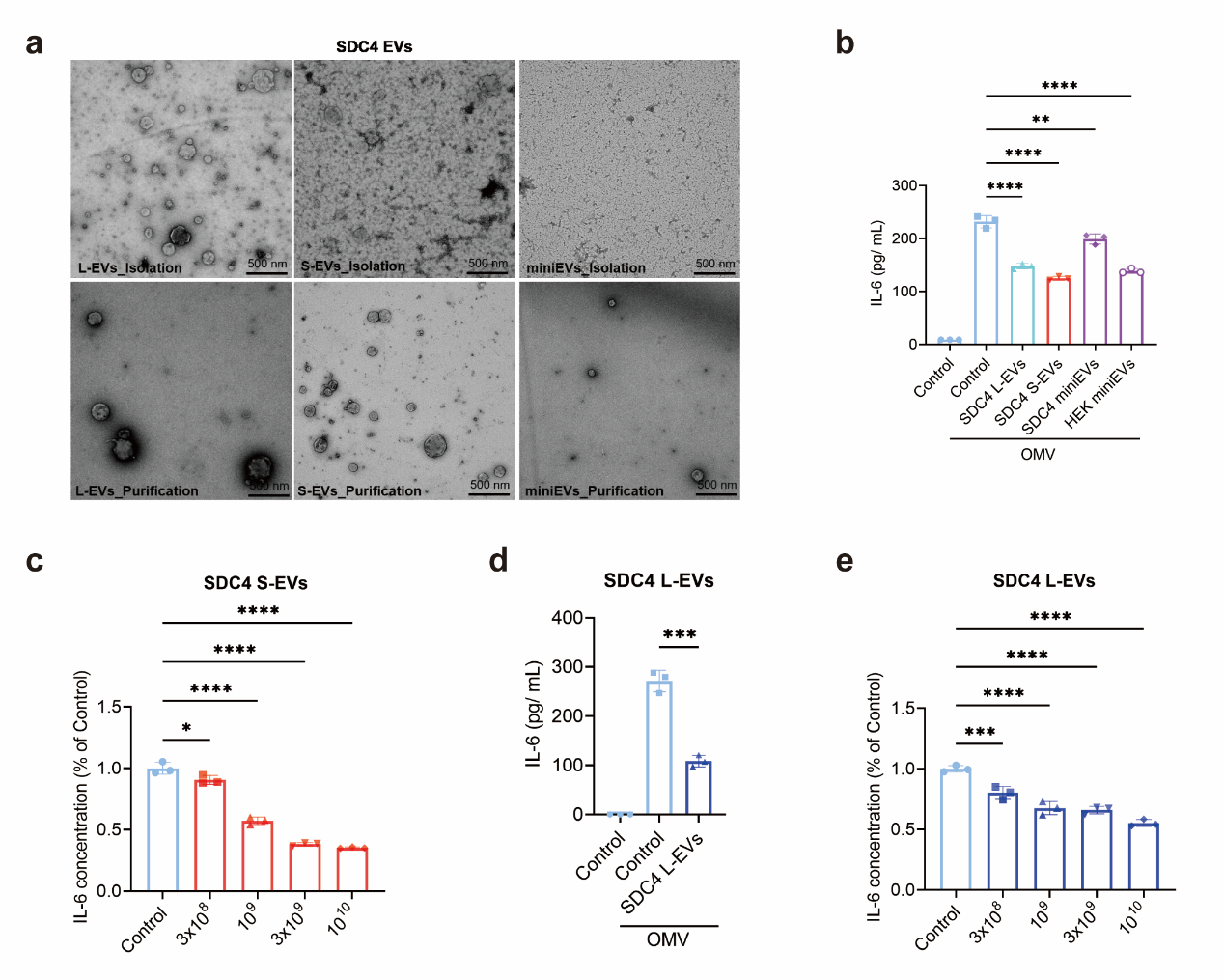
**Extended Figure 4. Characterization and anti-inflammation function of L-EVs, S-EVs and miniEVs of SDC4 overexpressing HEK293F cells.** **a)** TEM images of SDC4 L-EVs, S-EVs, and miniEVs after ultracentrifugation (called “isolation” – upper panels) and after iodixanol density gradient (called “purification” – lower panels). **b)** IL-6 concentration in the supernatant of RAW 264.7 cells after exposure to OMV (100 ng/mL) followed by treatment with EVs (10^9^ /mL). **c)** IL-6 concentration in the supernatant of RAW 264.7 cells after exposure to OMV (100 ng/mL) followed by treatment of increasing concentrations of SDC4 S-EVs. **d)** IL-6 concentration in the supernatant of RAW 264.7 cells after exposure to OMV (100 ng/mL) followed by treatment with SDC4 L-EVs _ isolation (10^9^ /mL). e. IL-6 concentration of the supernatant of RAW264.7 cells after exposure to OMV (100 ng/mL) followed by treatment of increasing concentrations of SDC4 L-EVs _ isolation. Data were analyzed using one-way ANOVA. *, *P* <0.05; **, *P* < 0.01; ***, *P* < 0.001; ****, *P* < 0.0001.

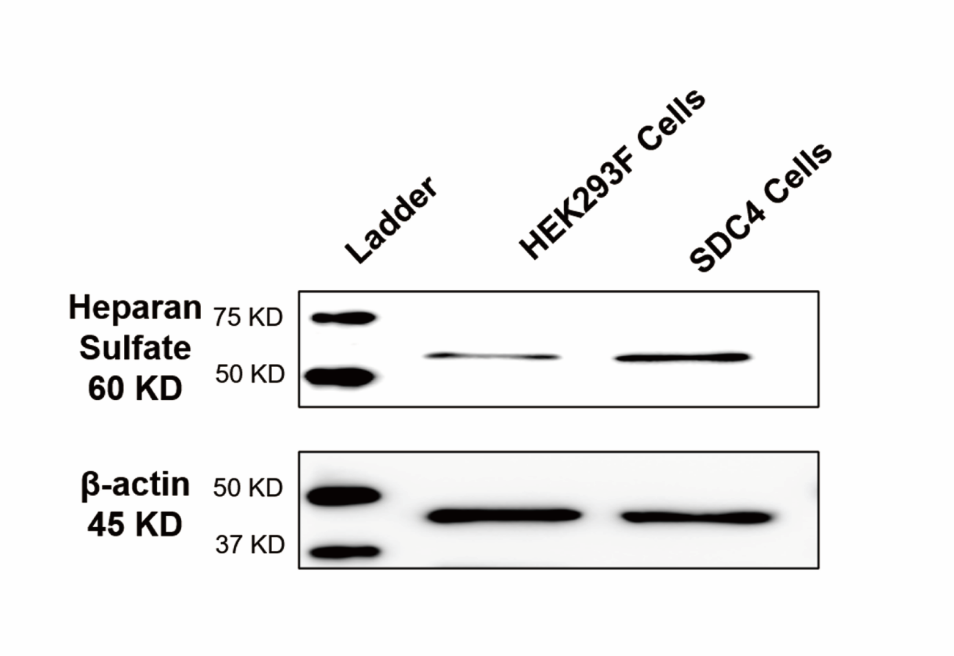

**Extended Figure 5 Western blot of heparan sulfate expression of wild-type HEK293F cells vs SDC4 overexpressing HEK293F cells.**
